# Oncogenic RAS sensitizes cells to drug-induced replication stress via transcriptional silencing of P53

**DOI:** 10.1101/2021.06.14.448289

**Authors:** Hendrika A. Segeren, Elsbeth A. van Liere, Frank M. Riemers, Alain de Bruin, Bart Westendorp

## Abstract

Cancer cells often experience high basal levels of DNA replication stress (RS), for example due to hyperactivation of oncoproteins like MYC or RAS. Therefore, cancer cells are considered to be sensitive to drugs that exacerbate the level of RS or block the intra S-phase checkpoint. Consequently, RS-inducing drugs including ATR and CHK1 inhibitors are used or evaluated as anti-cancer therapies. However, drug resistance and lack of biomarkers predicting therapeutic efficacy limit efficient use. This raises the question what determines sensitivity of individual cancer cells to RS. Here, we report that oncogenic RAS does not only enhance the sensitivity to ATR/CHK1 inhibitors by directly causing RS. Instead, we observed that HRAS^G12V^ dampens the activation of the P53-dependent transcriptional response to drug-induced RS, which in turn confers sensitivity to RS. We demonstrate that inducible expression of HRAS^G12V^ sensitized retina pigment epithelial (RPE-hTERT) as well as osteosarcoma (U2OS) cells to ATR and CHK1 inhibitors. Using RNA-sequencing of FACS-sorted cells we discovered that P53 signaling is the sole transcriptional response to RS. However, oncogenic RAS attenuates the transcription of P53 and its target genes. Accordingly, live cell imaging showed that HRAS^G12V^ exacerbates RS in S/G2-phase, which could be rescued by stabilization of P53. Thus, our results demonstrate that transcriptional control of P53 is a prime determinant in the response to ATR/CHK1 inhibitors and show that hyperactivation of the MAPK pathway impedes this response. Our findings suggest that the level of oncogenic MAPK signaling could predict sensitivity to intra-S-phase inhibition in cancers with intact P53.

## Introduction

Preservation of genomic integrity is essential for life and therefore strictly controlled by DNA damage checkpoints. However, excessive activity of oncoproteins such as MYC, RAS and Cyclin E pose a threat to faithful propagation of DNA by induction of replication stress (RS) (1,2). RS is referred to as impediments during DNA replication and thus includes stalling and collapsing of replication forks. Emerging evidence uncovers RS in the vast majority of cancer cells and is therefore considered a hallmark of cancer (3).

Activation of the intra S-phase checkpoint is the key response to RS and orchestrated by two central kinases, Ataxia Telangiectasia and Rad3-related protein (ATR) and Checkpoint Kinase 1 (CHK1) (4). Since high levels of RS are detrimental for survival, cancer cells heavily rely on the intra S-phase checkpoint (5–7). Accordingly, an extra allele of CHK1 protects against oncogene-induced RS (8) and high levels of CHK1 confer resistance to RS-inducing drugs (9). The dependency of cancer cells on the intra S-phase checkpoint has led to the development of ATR and CHK1 inhibitors (reviewed in (10)). These inhibitors are combined with classical chemotherapeutic drugs to completely exhaust the intra S-phase checkpoint and together referred to as RS-inducing drugs. However, the lack of knowledge on parameters that predict sensitivity to these drugs limits efficient clinical use, while drug resistance remains a major problem (11). In contrast, selection of BRCA mutant tumors for PARP inhibitor treatment shows that patient selection can dramatically improve drug efficiency. Interestingly, the lack of P53 can sensitize cells to ATR and CHK1 inhibition (12), suggesting a potential selection criterium for treatment with RS-inducing drugs. Nonetheless, P53 mutation status failed to predict the response to inhibitors of the intra S-phase checkpoint in xenograft experiments and clinical trials (13–15).

Besides P53, little focus has been paid to the role of oncogenic alterations in the response to RS-inducing drugs. Hyperactivation of the MAPK pathway by mutations in one of the RAS isoforms is one of the most frequent alterations in human malignancies (16). Oncogenic RAS elevates ROS and increases global transcription rates leading to high basal levels of RS (17,18). As a result, RAS-mutant cells heavily rely on the intra S-phase checkpoint for survival (5,7) and tumors with oncogenic RAS are promising candidates for treatment with RS-inducing drugs. Interestingly, conflicting studies showed that RAS could either increase or reduce P53 levels (19,20). More recently, ERK signaling, downstream of RAS, was shown to moderate the pulsatile behavior of P53 signaling, which permits proliferation in the presence of mild damage (21). However, despite the key role of ERK signaling in the proliferation-quiescence decision (22), the effect of oncogenic RAS on cell fate decisions upon intra-S-phase checkpoint inhibition has been neglected.

In addition to the variety of mutations a tumor can harbor, cell-intrinsic heterogeneity in cell cycle phase and level of RS blur the picture when evaluating the response to RS-inducing drugs. Therefore, it is of critical importance to firstly determine how a single cell responds to RS-inducing drugs and secondly evaluate how mutations affect this process to optimize anti-cancer treatment with RS-inducing drugs.

Here we use non-transformed human cells containing fluorescent cell cycle and RS reporters to show that transcriptional regulation of P53 is essential to control the response to RS. We demonstrate that inducible HRAS^G12V^ impinges the RS response by transcriptional downregulation of P53, resulting in downregulation of DNA repair genes in S/G2 phase. Suppression of the P53-dependent gene transcription program by oncogenic HRAS sensitizes cancer cells to RS-inducing drugs, and decreases long-term viability after transient treatment with these drugs.

## Results

### *Development of an* in vitro *model with multiple reporters to study the RS-response in living single cells*

The response of cancer cells to RS is most likely affected by the combined action of multiple oncogenic changes. This makes it complicated to address the effect of single mutations. We therefore utilized human non-transformed hTERT-RPE1 cells in which we introduced oncogenic HRAS (HRAS^G12V^) under the control of a Tet repressor. Upon administration of doxycycline, these cells hyperactivated the MAPK signaling pathway (Figure S1A/B). In accordance with this oncogenic change, HRAS^G12V^ stimulated cell proliferation (Figure 1A). Remarkably, induction of HRAS^G12V^ was sufficient to allow transformation and subcutaneous tumor growth of RPE cells in mice (Figure 1B).

**Figure 1:**
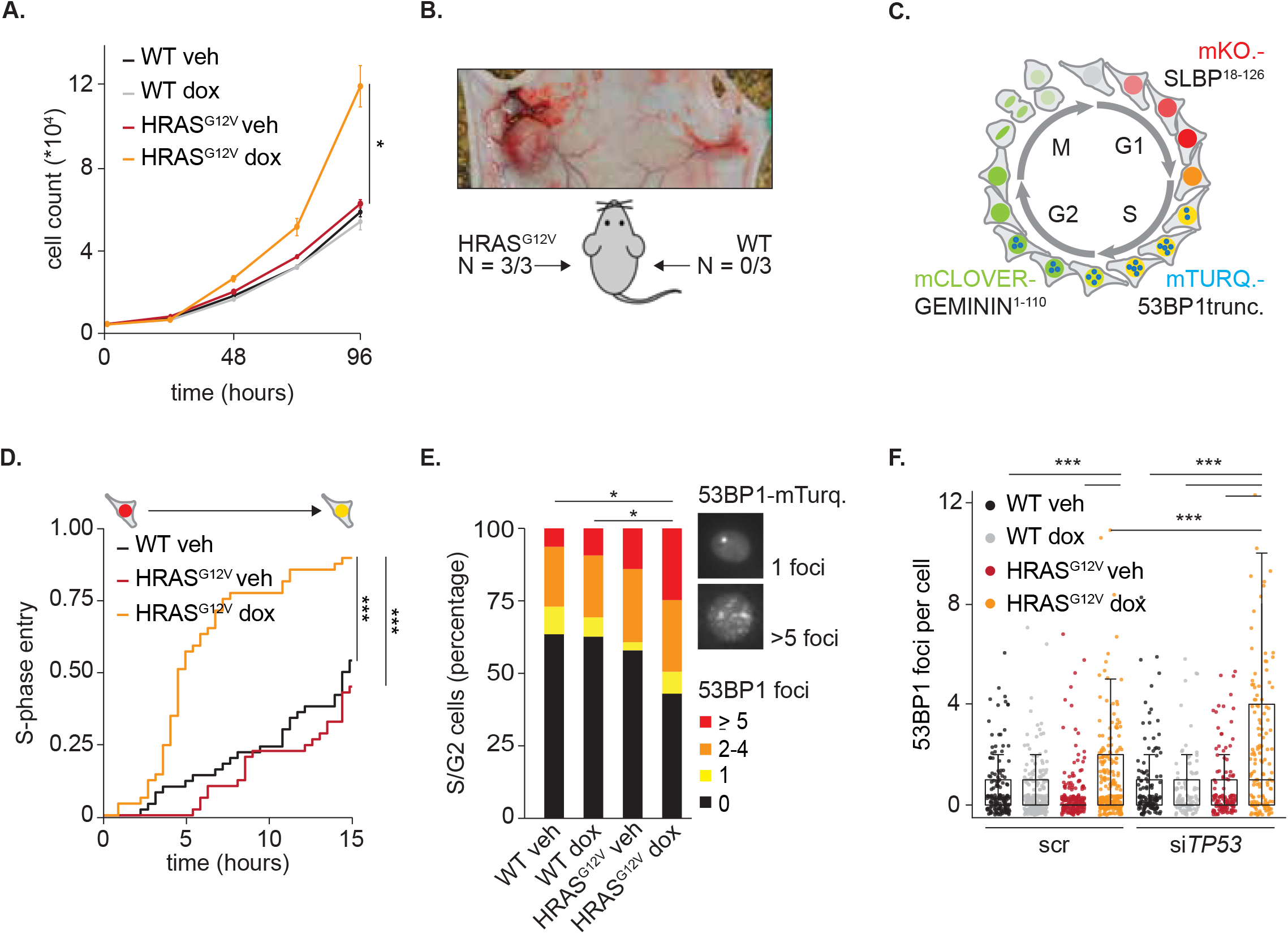
Inducible HRAS^G12V^ shortens G1-phase and induces mild RS. **A** Cell proliferation, as measured by cell counting, of RPE WT or HRAS^G12V^ cells with or without doxycycline. Measurement indicates mean ± s.e.m., statistical analysis was performed with a Kruskal-Wallis test. **B** Representative image and quantification of RPE WT and HRAS^G12V^ cells injected in mice. This resulted in tumor formation in cells expressing HRAS^G12V^ (left). **C** Schematic representation of FUCCI4 system and fluorescent tagged 53BP1 during cell cycle progression. **D** Cumulative frequency plot of WT and HRAS^G12V^ cells with or without doxycycline which were in G1 phase at the start of imaging. The time until S-phase entry of at least 25 individual cells per condition was measured. Differences were statistically evaluated with a Log-rank test. **E** Bar chart showing the number of 53BP1 foci, as read out for RS, per S/G2-phase cell in the absence or presence of 24 hours oncogenic RAS induction. At least 100 cells per condition were analyzed. Statistical differences were evaluated with a Chi-Square test corrected for multiple comparisons. **F** Dot plot showing the number of 53BP1 foci in individual cells 24 hours after siRNA and doxycycline treatment for indicated conditions. Per condition at least 100 cells were evaluated. Differences were statistically tested using a Kruskal-Wallis test with post-hoc Dunnett’s test with Benjamin Hochberg correction.

We then investigated which phase of the cell cycle was accelerated by HRAS^G12V^. Therefore, we introduced the Fluorescent Ubiquitination-based Cell Cycle Indicator (FUCCI) 4 system (23) into these cells (Figure 1C). Live cell imaging revealed that the increased proliferation speed of cells with HRAS^G12V^ could be attributed to a shorter G1-phase, while length of other phases was unaffected (Figure 1D/S1C).

G1-phase shortening and premature S-phase entry can cause RS (24). We evaluated the effect of oncogenic RAS signaling on DNA replication using DNA fiber assays. In this assay cells are pulsed with fluorescent labelled nucleotides for a set amount of time, from which origin firing and replication speed can be inferred. Only a slight increase in the number of fired origins (Figure S2A), and no effect on DNA replication speed was observed (Figure S2B). Moreover, in whole cell lysates no abundant CHK1 phosphorylation was present, indicating inactive CHK1 and the absence of severe RS (Figure S2C). However, also mild RS can induce DNA damage (25). Indeed, we observed a small but significant increase in γH2AX foci in cells expressing HRAS^G12V^ (Figure S2D).

To in more detail evaluate the effect of oncogenic RAS on DNA replication in living single cells, we incorporated a fluorescent tagged truncated version of 53BP1 in FUCCI4-expressing cells (Figure 1C). The formation of 53BP1 foci is indicative of RS-induced DNA damage (26). In line with the previous observations, we saw a small but significant increase in 53BP1 foci per cell in the presence of HRAS^G12V^ in S/G2 phase (Figure 1E). Although early reports showed that oncogenic RAS induces RS resulting in a P53 mediated growth arrest (Di Micco et al., 2006), we found that expression of HRAS^G12V^ in RPE1-hTERT cells did not inhibit proliferation. This was presumably due to the presence of telomerase (27) and intact cell cycle checkpoints. As P53 fulfills a pivotal role in protection against RS (28,29), we investigated if intact P53 was required to limit endogenous DNA damage in these cells, by knocking down P53 with RNAi. Remarkably, knockdown of *TP53* increased the number of 53BP1 foci in HRAS^G12V^ expressing cells, indicating more RS (Figure 1F). Collectively, these data show that acute induction of HRAS^G12V^ in unperturbed RPE cells causes surprisingly mild RS as detected by 53BP1 foci, despite dramatic G1-phase shortening and unleashing malignant transformation. However, P53 is required to minimize DNA damage and maintain genomic integrity in HRAS^G12V^ expressing cells.

### Oncogenic RAS sensitizes cells to RS-inducing drugs

Excessive RS is deleterious for cell survival, thus cancer cells are predicted to heavily rely on the intra S-phase checkpoint to control RS. Since induction of HRAS^G12V^ causes mild RS, we asked whether this would exacerbate the level of RS induced by inhibitors of the intra-S-phase checkpoint, of which ATR and CHK1 are the main players. To fully exhaust this checkpoint and induce severe RS, the ATR inhibitor Ceralasertib (ATRi) or CHK1 inhibitor Prexasertib (CHK1i) can be combined with a low dose of drugs which interfere with DNA replication such as the nucleoside analogue gemcitabine, a strategy also evaluated in clinical trials (Baillie, 2020). We first confirmed in normal RPE cells that CHK1i and gemcitabine caused RS and DNA damage in a synergistic manner, as seen by phosphorylation of CHK1 on Serine 345 and γH2AX staining (Figure 2A/B). A dose of 10 nM CHK1i efficiently inhibited CHK1 activation, as shown by the absence of its autophosphorylation on Serine 296 (Figure 2A). The combination therapy of CHK1i and gemcitabine blocked DNA replication and cell proliferation independent of HRAS^G12V^ status (Figure S3A/B). However, after 48 hours of treatment the level of DNA-damage was strongly elevated in cells overexpressing HRAS^G12V^ as measured by the number of 53BP1 foci per cell (Figure 2C). Using an ATRi instead of CHK1i had a similar effect (Figure 2D), indicating that the higher sensitivity of HRAS^G12V^ cells is not drug-specific, but a consequence of intra S-phase checkpoint inhibition. To prove that this increased sensitivity after HRAS^G12V^ induction was not cell line specific, we repeated the experiments in U2OS osteosarcoma cells. Indeed, U2OS cells also present mild RS upon expression of HRAS^G12V^, and were more sensitive to CHK1i + gemcitabine treatment (Figure S3C/D/E). Because these RS-inducing drugs will not be given continuously to patients, we analyzed recovery of the cells after drug withdrawal. Colony formation assays showed that recovery from 48 hours of CHK1i + gemcitabine was significantly impaired in RPE cells expressing oncogenic HRAS compared to non-transformed RPE cells (Figure 2E/F/G). In contrast, colony formation capacity under unperturbed conditions was not affected by HRAS^G12V^ induction (Figure S3F). Together, these data show that oncogenic RAS sensitizes cells to RS-inducing drugs, as seen by elevated DNA damage and impaired recovery after drug withdrawal.

**Figure 2:**
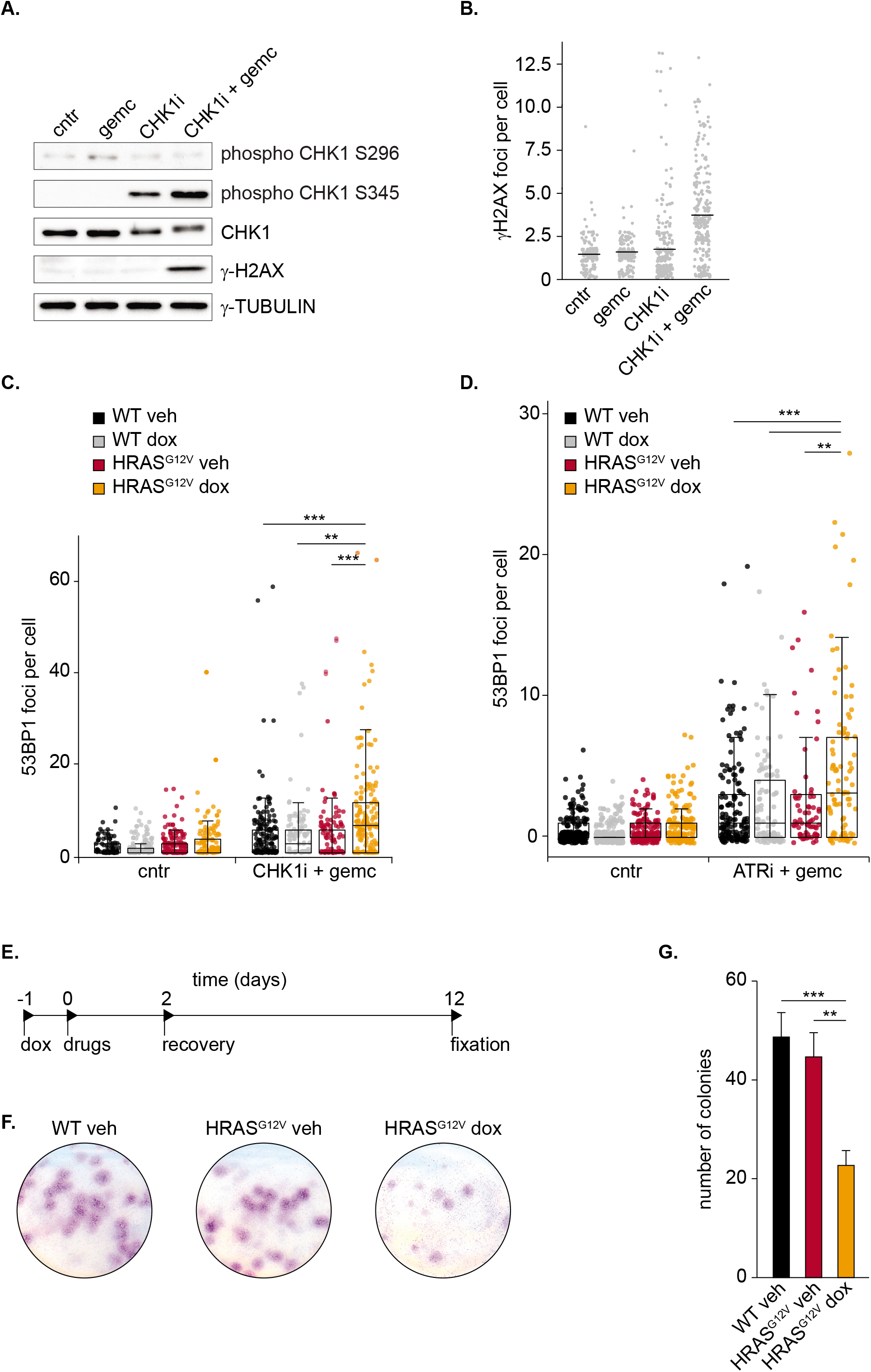
Oncogenic RAS sensitizes cells to replication stress-inducing drugs. **A** Immunoblot showing phosphorylation of CHK1 and γH2AX in RPE WT cells 24 hours after indicated drug treatment. **B** Quantification of γH2AX immunofluorescence staining of RPE WT cells 24 hours after indicated drug treatment. **C** Dot plot showing 53BP1 foci in individual RPE WT or HRAS^G12V^ cells before and 48 hours after treatment with CHK1i + gemcitabine. At least 50 cells per condition were analyzed. Differences were statistically tested using a Kruskal-Wallis test with post-hoc Dunnett’s test with Benjamin Hochberg correction. **D** Same as C, but with ATRi + gemcitabine. **E** Experimental setup of colony formation assay. Cells were fixed and stained 10 days after drug (CHK1i + gemcitabine) washout and replating. **F** Representative pictures of a colony formation assay described in E. **G** Quantification of three colony formation assays. Bars represent mean (sum of 3 technical replicates, mean of 3 independent experiments) ± s.e.m., statistical analysis was performed with a Kruskal-Wallis test.

### P53 signaling is essential and responsible for the response to RS

Repetitive treatment of cells which initially recovered from treatment with RS-inducing drugs yielded similar recovery rates (Figure 3A). Therefore, we hypothesized that the reduced outgrowth of HRAS^G12V^ cells compared to wild-type cells is caused by a different transient response to drug treatment and not by permanent resistant or hypersensitive subpopulations of cells. To explore gene expression programs that could explain the higher sensitivity of HRAS^G12V^-expressing cells, we performed RNA-sequencing of S-phase cells with and without HRAS^G12V^ before and 16 hours after treatment with CHK1i + gemcitabine. We FACS-sorted S-phase cells using the FUCCI4 reporters to avoid bias from differences in cell cycle phase distributions (Figure S4A). First, we evaluated the effect of overexpression of HRAS^G12V^ alone. Although induction of oncogenic RAS yielded robust transcriptional changes, they were not related to RS (Figure S4B). We then evaluated the effect of RS-inducing drugs. In wild-type RPE cells we identified just under 100 up- and 30 downregulated genes during drug treatment in S-phase (Figure S4C). Gene ontology analysis revealed that the majority of the upregulated genes is involved in P53 signaling (Figure 3B). A subset of these P53 target genes, albeit at a lower magnitude, was also increased by CHK1i + gemcitabine treatment in HRAS^G12V^-expressing cells (Figure 3B, S4C). Additionally, in accordance with previously published results, TGF-beta signaling was hampered in cells with oncogenic RAS (30). Furthermore, differential expression analysis showed that activation of the P53 program is the prime response to RS in RPE cells with and without HRAS^G12V^ overexpression (Figure 3C).

**Figure 3:**
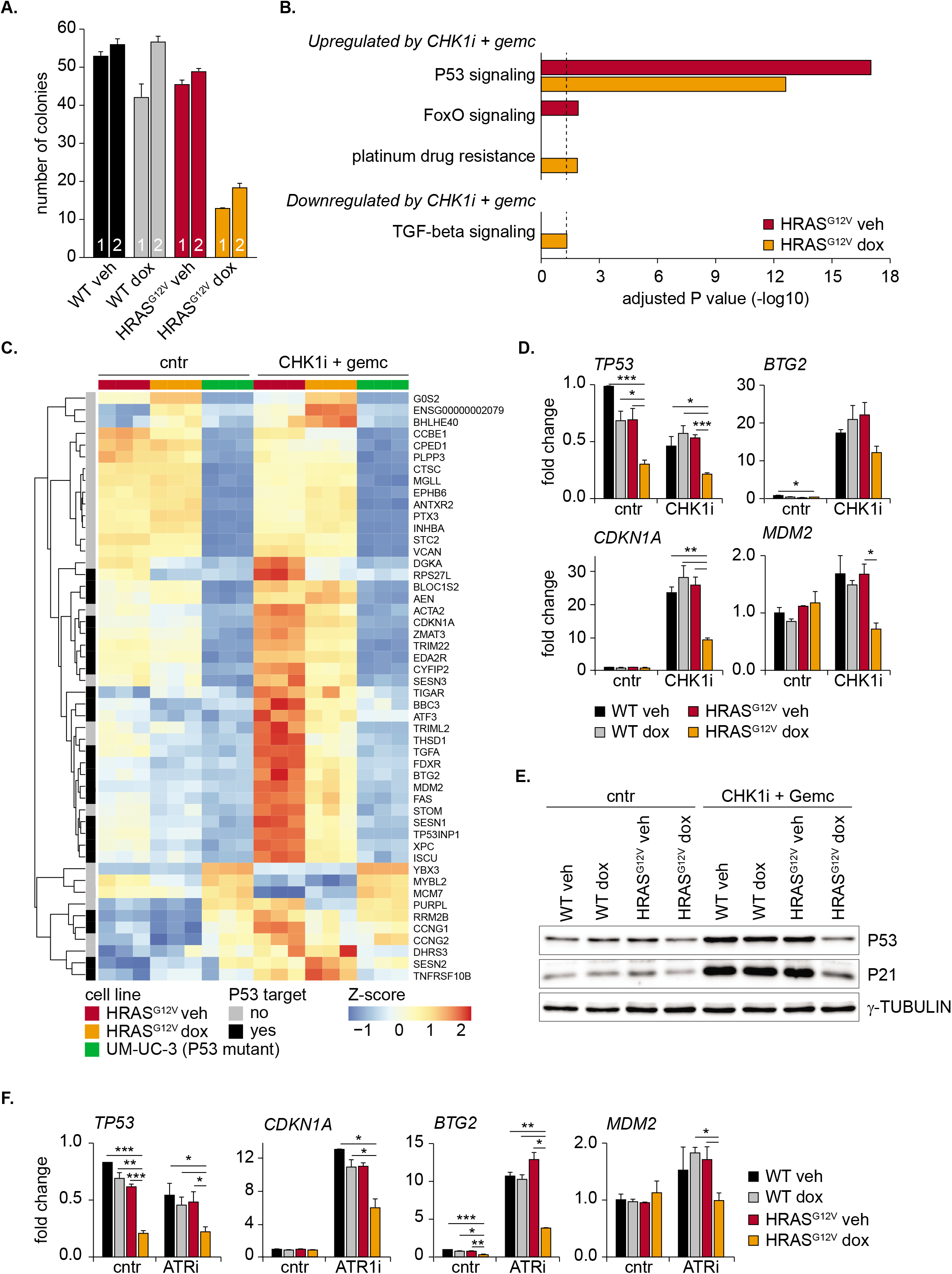
P53 signaling is essential and responsible for the response to replication stress. **A** Quantification of colony formation assay of indicated conditions after 48 hours of treatment with CHK1i + gemcitabine (1) or after 2 rounds of treatment separated by a 10-day recovery period (2). Colonies were quantified 10 days after end of treatment. Bars represent mean ± s.e.m. **B** Pathway analysis of differentially expressed genes in HRAS^G12V^ S-phase cells before and 16 hours after treatment with CHK1i + gemcitabine. Significantly changed genes with a fold change of at least 1.5 were selected for analysis. Black dotted line indicates *P* value of 0.05. **C** Heatmap of the 50 most significantly changed genes in wild-type S-phase RPE cells before and 16 hours after treatment with CHK1i + gemcitabine. The normalized expression of these genes in S-phase RPE cells with and without HRAS^G12V^ and UM-UC-3 S/G2-phase cells is shown. P53 target genes identified using meta-analysis are indicated (62). **D** Quantitative PCR showing reduced expression of *TP53* and P53 target genes in HRAS^G12V^ cells before and after treatment with CHK1i + gemcitabine. Bars represent mean ± s.e.m. of 2 independent experiments. Statistical differences were evaluated using a Kruskal-Wallis test. **E** Immunoblot showing reduced P53 and P21 levels in cells with oncogenic RAS 24 hours after doxycycline treatment in the absence and presence of CHK1i + gemcitabine. **F** Same as D, but now with ATRi + gemcitabine.

To identify potential P53-independent RS response pathways, we also included the P53 deficient UM-UC-3 bladder cancer cell line before and after treatment with RS-inducing drugs in our RNA-sequencing analysis. The P53-dependent response was completely absent in these cells, and remarkably, only five differentially expressed genes could be identified, indicating that the transcriptional changes after CHK1i + gemcitabine treatment are entirely depending on P53 (Figure 3C, S4C). As a result, UM-UC-3 cells were hypersensitive, as shown by severely impaired proliferation rates, to treatment with a CHK1i + gemcitabine (Figure S4D). Our data show that transcriptional changes after intra S-phase checkpoint inhibition are P53-dependent, and underscore that P53 is essential and responsible for an efficient response to RS.

### RAS mediated downregulation of P53 transcripts compromises the RS response in G2 phase

The data presented above underscores the importance of P53 in response to RS. Nonetheless, this P53 response upon treatment with the CHK1i + gemcitabine was dampened in cells expressing HRAS^G12V^ compared to wild-type cells (Figure 3C). P53 activity is known to be predominantly regulated by the stability of the P53 protein. But quantitative PCR showed that *TP53* transcripts were strongly downregulated in cells with oncogenic RAS under both unperturbed conditions and during CHK1i + gemcitabine treatment (Figure 3D). This resulted also in lower P53 protein levels (Figure 3E). Subsequently, prime P53 targets were downregulated in these cells (Figure 3D). Similarly, attenuated expression of *TP53* and its targets was observed after treatment of HRAS^G12V^ expressing cells with an ATR inhibitor (Figure 3F). Moreover, U2OS cells with inducible HRAS^G12V^ exhibited reduced transcript levels of *TP53* (Figure S4E). Thus, the downregulation of *TP53* by oncogenic RAS is a general response to RS-inducing drugs and is independent of cell type.

Next, we sought to determine how oncogenic RAS signaling can transcriptionally downregulate P53. A likely candidate is the co-transcriptional regulator RAS Responsive Element Binding Protein 1 (RREB1). Oncogenic RAS phosphorylates RREB1 which cooperates with SMAD proteins to control transcription of SMAD target genes (31). Interestingly, RREB1 is also shown to bind and activate the P53 promotor region (32), linking oncogenic RAS signaling to P53. However, knock down of *RREB1* did not decrease *TP53* transcript levels and its downstream targets (Figure S5A). We then exploited our RNA-sequencing dataset to identify putative transcriptional regulators of P53 that were affected by HRAS induction. We selected genes with at least a 1.5-fold change in expression comparing wild-type and HRAS^G12V^ cells, DNA binding capacity and a previously described link with P53 expression. This resulted in a list of four candidates, two potential repressors (CEBP-beta and KLF4) and two potential activators (SMAD3 and KLF9) which are up- and downregulated upon HRAS^G12V^ expression respectively (Figure S5B). We evaluated if siRNA oligos targeting these candidates could rescue or mimic the *TP53* levels in cells with HRAS^G12V^. Nevertheless, despite efficient depletion of the putative regulators, no rescue or phenocopy of the effect of HRAS^G12V^ on P53 was observed (Figure S5C-G). Thus, HRAS^G12V^ activation does not directly control *TP53* expression via one of the selected transcription regulators, although more elaborate studies using proteomics or screening approaches may unveil single regulators connecting oncogenic RAS to *TP53* transcription.

The general dampened P53 response in cells with HRAS^G12V^ was identified using RNA-sequencing of S-phase cells. However, P53 regulates a plethora of genes involved in cell cycle progression, apoptosis and DNA repair whose function is not limited to S-phase. We therefore analyzed the expression of a subset of P53 target genes with different functions, before and after treatment with RS-inducing drugs and in different cell cycle phases (Figure S4A). Although downregulation of *TP53* was evident in all cell cycle phases (Figure 4A), most abundant downregulation of target genes was present in cells residing in G2 phase (Figure 4B). This included genes related to DNA repair such as *XPC, PCNA* and *RRM2B.* These data show that oncogenic RAS blocks the expression of P53 and its target genes during the RS response, resulting in strong downregulation of multiple P53-responsive DNA repair genes and cell cycle inhibitors.

**Figure 4:**
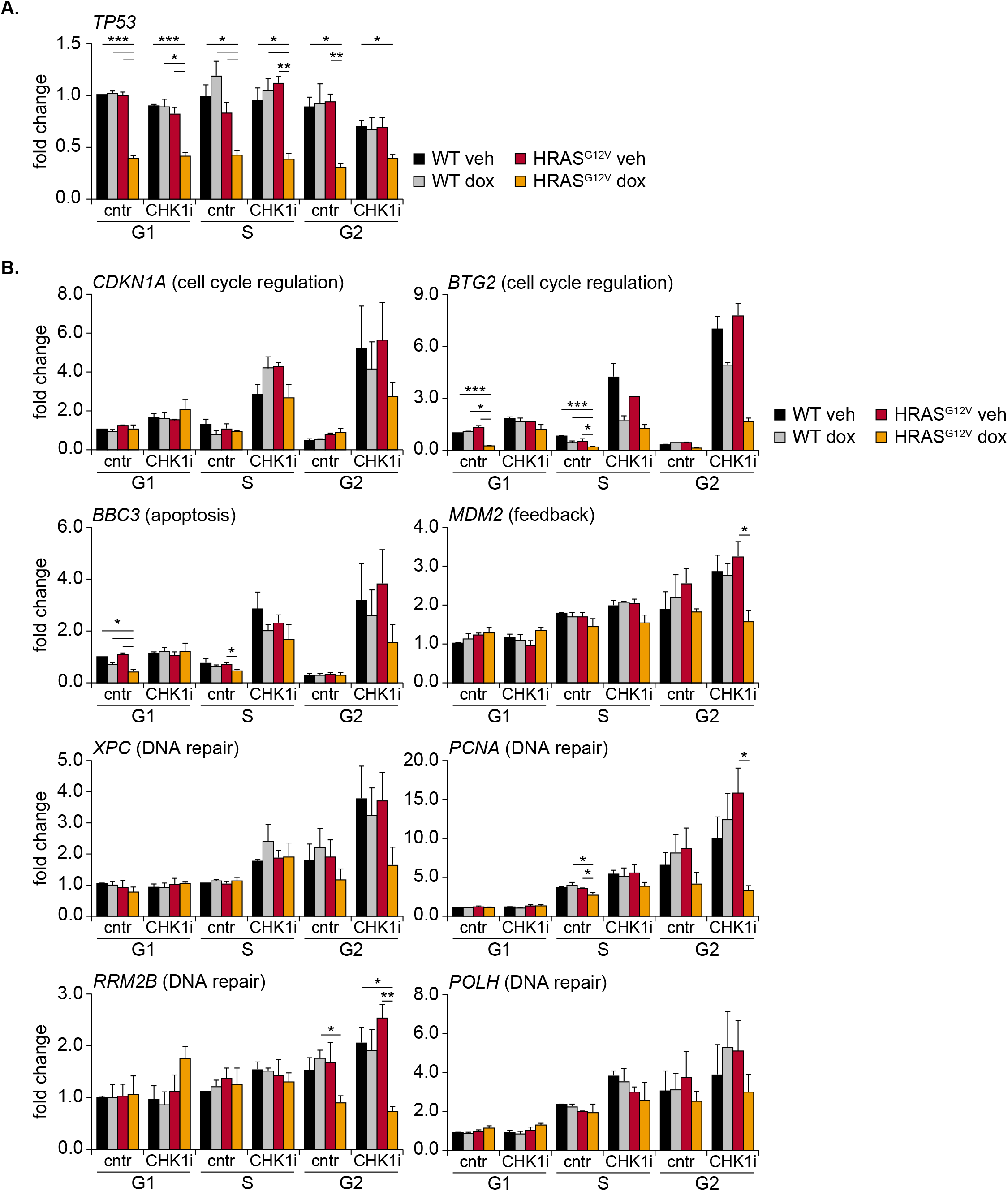
Oncogenic RAS compromises the RS response in G2 phase. **A** Quantitative PCR showing downregulation of *TP53* in HRAS^G12V^ dox cells independent of cell cycle phase. Bars represent mean ± s.e.m. of 3 independent experiments. Statistical differences were evaluated using a Kruskal-Wallis test. **B** Quantitative PCR of indicated P53 target genes in different cell cycle phases. Bars represent mean ± s.e.m. of 3 independent experiments. Statistical differences were evaluated using a Kruskal-Wallis test.

### HRAS^G12V^ exacerbates level of RS and delays cell cycle exit during drug treatment

Since P53 is a critical regulator of cell cycle progression and DNA repair, the dampened P53 response during S/G2-phase in cells with oncogenic RAS, can be expected to have profound impact on cell cycle fates. To evaluate this at a single cell level, we performed a live cell imaging experiment. We treated cells expressing the FUCCI4 markers as well as truncated 53BP1, to monitor DNA damage, with a CHK1i + gemcitabine and followed the fates of individual cells in response to treatment (Figure 5A/B). We did not observe cells undergoing cell death, possibly due to the presence of intact cell cycle checkpoints in hTERT-RPE1 cells. To evaluate other cell fates, single cell traces (Figure 5B) were combined in heatmaps, and ordered according to the moment of the first mitosis (Figure 5C-F). These heatmaps confirmed that cells with HRAS^G12V^ experience elevated levels of endogenous RS, as measured by 53BP1 foci during S/G2-phase, which was exacerbated upon treatment with RS-inducing drugs (Figure 5C-F/6A). Although CHK1i + gemcitabine treatment initially decreased the average number of 53BP1 foci throughout S-phase in all cell lines, elevated levels of RS-induced DNA damage in cells with oncogenic RAS were clearly evident during G2-phase (Figure 6A). Offspring of cells that were able to undergo mitosis showed a low number of 53BP1 foci in the next G1 (Figure 6A). These foci in G1 cells, usually referred to as nuclear bodies, indicate unresolved DNA damage, but did not differ substantially between any of the conditions (25).

**Figure 5:**
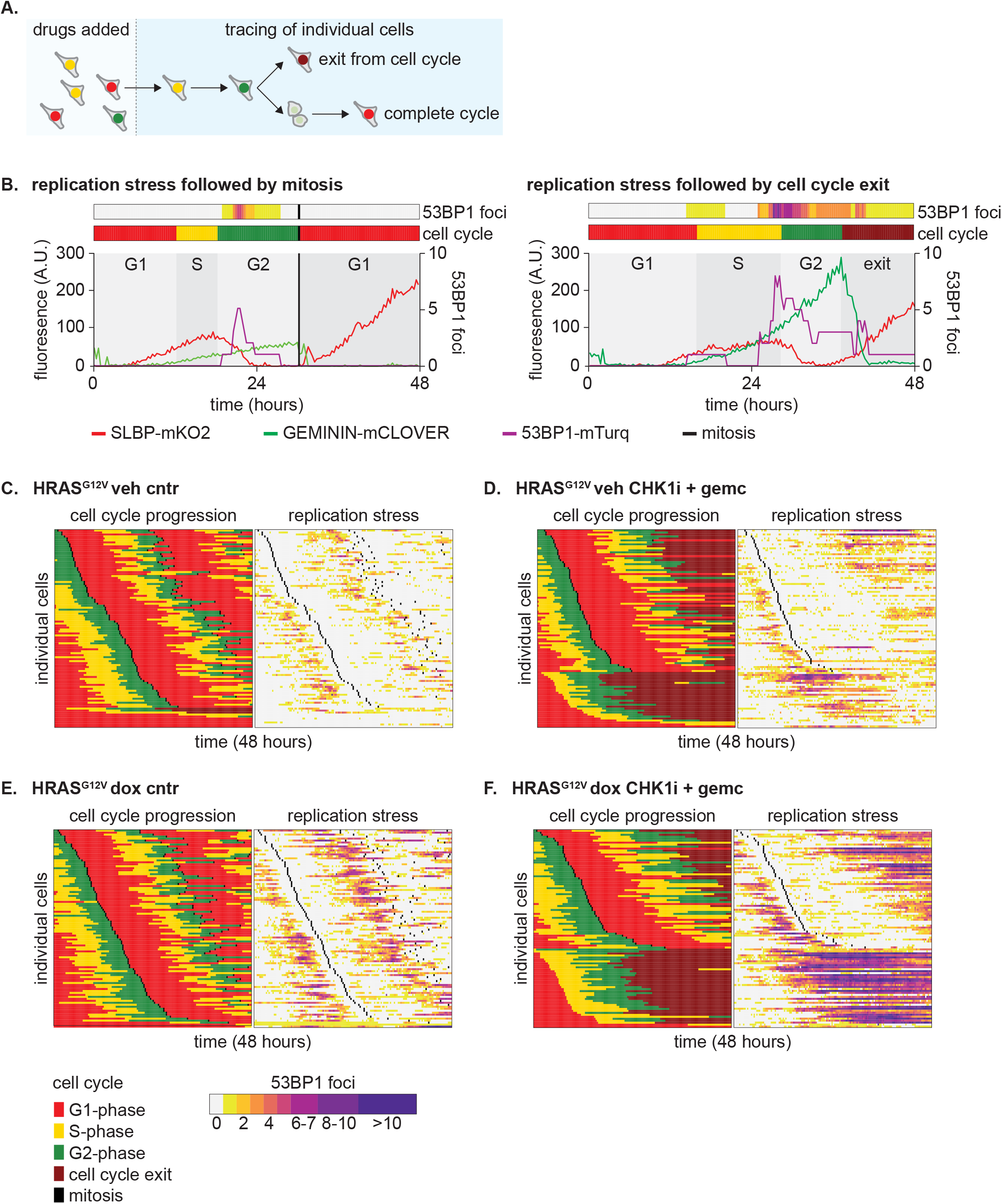
Single cell analysis of the replication stress response. **A** Schematic representation of cell tracing strategy and potential fates in live cell imaging experiments. **B** Representative cell traces of two cells showing 53BP1 foci and cell cycle progression in the presence of CHK1i + gemcitabine. The left cell completes mitosis, the right cell exits the cell cycle without mitosis. **C** Heatmap showing cell cycle progression and 53BP1 foci in 100 individual HRAS^G12V^ veh cells under control conditions. Each row represents a single cell. Cells were order based on the moment of first mitotic entry. **D** Same as C, but now cells were treated with CHK1i + gemcitabine. **E** Same as C, but now HRAS^G12V^ dox cells. **F** Same as C, but now HRAS^G12V^ dox cells that were treated with CHK1i + gemcitabine.

**Figure 6:**
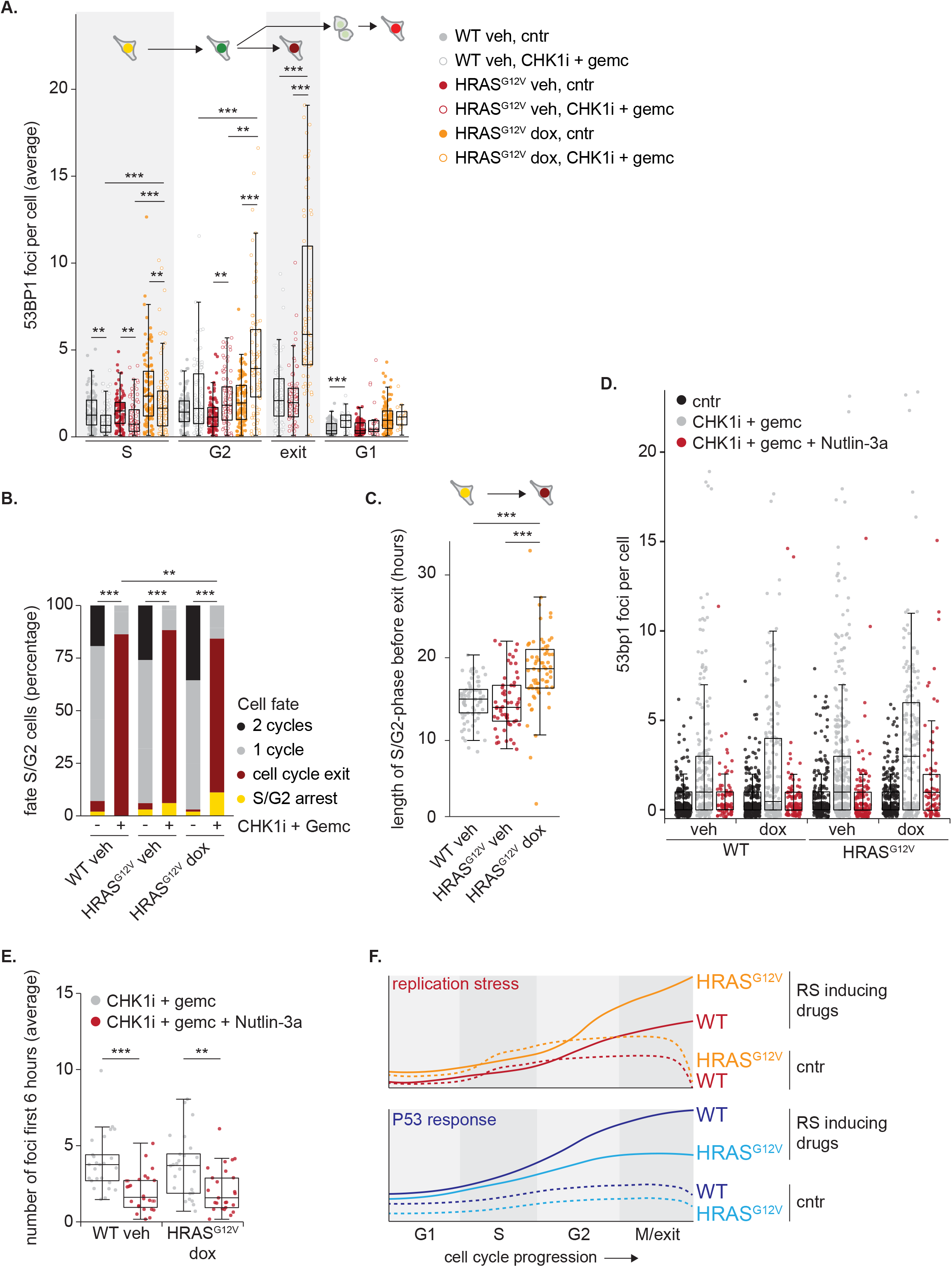
P53 silencing by HRAS^G12V^ exacerbates RS during drug treatment. **A** Quantification of live cell imaging data showing the average number of 53BP1 foci per image per cell in different cell cycle phases. Cells were included from S-phase entry onwards. 100 cells were analyzed, and statistical analysis was performed with a Kruskal-Wallis test with post-hoc Dunnett’s test with Benjamin Hochberg correction. **B** Bar chart showing the frequencies of different cell fates of S-phase cells after treatment with CHK1i + gemcitabine. 100 cells per condition were evaluated and statistical difference was tested with a Chisquare test corrected for multiple comparisons. **C** Dot plot in which the duration from S-phase entry until cell cycle exit is shown. Only cells which enter S phase and subsequently exit the cell cycle upon treatment with CHK1i + gemcitabine were included. Statistical differences were evaluated using a Kruskal-Wallis test. **D** Quantification of 53BP1 foci per cell, cultured for 48 hours in the presence or absence of CHK1i + gemcitabine combined with Nutlin-3a. **E** Quantification of the live cell imaging data shown in supplemental Figure 6C/D. Average number of 53BP1 foci in S/G2-phase cells during the first 6 hours of treatment were plotted. 25 cells per condition were analyzed. Statistical differences were evaluated using a Kruskal-Wallis test and post-hoc Dunnett’s test. **F** Schematic overview representing the key findings of the current study. Oncogenic RAS induces endogenous RS which can be exacerbated by treatment with RS-inducing drugs. Activation of the P53 transcription program is the prime response to RS. This response is compromised in cells with HRAS^G12V^ due to transcriptional downregulation of P53.

Although some cells completed mitosis in the presence of RS-inducing drugs, the majority exited the cell cycle by premature activation of APC/C^CDH1^, as seen by the loss of Geminin without mitosis (Figure 5B-F/6B). We asked if cells which exit the cell cycle in this manner experienced the highest levels of RS. However, when we classified cells according to cell cycle fate and analyzed the number of foci in the initial S-phase, we observed that cell cycle exit was not necessarily preceded by an S-phase with the highest levels of RS-induced DNA damage (Figure S6A). Instead, 53BP1 foci accumulate after exit from the cell cycle, specifically in cells with oncogenic RAS (Figure 5C-F/6A). This marked increase in 53BP1 foci in HRAS^G12V^ expressing cells exiting the cell cycle most likely represents a failure to repair DNA breaks (33).

Despite elevated levels of RS-induced DNA damage in G2-phase cells with oncogenic RAS compared to wild type cells, the cell cycle exit in CHK1i + gemcitabine treated cells after completion of S-phase was delayed by HRAS^G12V^ expression (Figure 6B/C). Potentially, the higher level of DNA damage in these cells has slowed down S-phase in a graded fashion (34), which could subsequently delay cell cycle exit. Additionally, P53-dependent P21 accumulation plays an important role in re-activating APC/C^CDH1^ to enforce the cell cycle exit. This P21 accumulation was impaired in cells expressing HRAS^G12V^ (Figure 4B). Thus, oncogenic RAS exacerbates the level of RS-induced DNA damage upon intra S-phase inhibition and delays exit from the cell cycle.

HRAS^G12V^ transcriptionally downregulates P53 resulting in attenuated activation of target genes which play a pivotal role in DNA repair. To bear out the hypothesis that a dampened, P53-induced, DNA-repair response directly underlies the increase in RS observed in cells with oncogenic RAS, we stabilized P53 using the MDM2 inhibitor Nutlin-3a. Nutlin-3a stabilized P53 target genes and resulted in marked decrease in 53BP1 foci formation after 48 hours of treatment with CHK1i + gemcitabine (Figure S6B/6D). Sustained P53 signaling is proven to affect cell fate and stimulate a permanent cell cycle arrest (35). Indeed, live cell imaging revealed that Nutlin-3a forced the vast majority of cells which were in early S-phase upon treatment with RS-inducing drugs to exit the cell cycle (Figure S6C/D). Despite this cell cycle exit, we observed that Nutlin-3a rescued the formation of 53BP1 foci, indicating less RS, during the first hours after the start of drug treatment (Figure 6E). Prolonged Nutlin-3a incubation in S-phase cells did not prevent the formation of 53BP1 foci (Figure S6C/D). Most likely, the continuous Nutlin 3a-induced accumulation of P53 leads to extremely high levels of its target P21 that completely blocks CDK activity and thereby impedes DNA replication and G2 progression (36).

Collectively, these data demonstrate that intra S-phase inhibition exacerbates RS in cells with oncogenic RAS via transcriptional downregulation of P53. Thus, our data predict that RAS mutations enhance sensitivity and impair recovery of cancer cells to treatment with RS-inducing drugs.

## Discussion

Oncogenes are described to confer sensitivity to ATR and CHK1 inhibition due to the presence of endogenous RS (5–7,37). We show that oncogenic RAS indeed exacerbates the level of RS upon treatment with inhibitors of the intra S-phase checkpoint. However, the mechanistic experiments presented here reveal that transcriptional downregulation of P53 by oncogenic RAS is a major determinant in sensitivity to RS-inducing drugs (Figure 6F). Although frequently mutated, P53 is intact in a substantial percentage of tumors harboring mutated RAS (38). The mutation status of P53 has been proposed to be predictive for sensitivity to ATR and CHK1 inhibitors, but clinical trials have not confirmed this yet (13). Thus, we propose that mechanisms controlling P53 activity such as RAS-dependent signaling must be taken into account when predicting sensitivity to intra-S-phase inhibitors in individual cancer patients. In line with this, our study provides a rationale to select cancer patients with hyperactivation of the MAPK pathway for treatment with intra S-phase checkpoint inhibitors. In contrast, therapies combining inhibitors of MAPK signaling and RS-inducing drugs should be carefully evaluated.

We detected attenuated P53 transcript levels in HRAS^G12V^ cells irrespective of cell cycle phase. Nonetheless, downregulation of target genes was mainly observed in S/G2-phase. Hence, it can be proposed that P53 executes its protective function during DNA replication, rather than by preventing cell proliferation (39). Indeed, prime P53 target genes involved in cell cycle arrest and apoptosis are dispensable for tumor suppression *in vivo* (40). This leaves energy metabolism, DNA repair, and preservation of genomic integrity as the key tumor-suppressive functions of P53. In line with a role in protecting DNA from damage, P53 is shown to prevent slowing and stalling of replication forks under conditions of genotoxic stress (29). This effect was at least in part dependent on the P53 target *MDM2,* which we also found downregulated in cells overexpressing oncogenic RAS (Figure 4C). In addition, we observed that the P53 target *POLH,* which encodes the translesion synthesis (TLS) polymerase η, was downregulated by HRAS^G12V^. TLS polymerases are required to facilitate DNA elongation over damaged DNA. Although this increases the risk of mutagenesis, it enhances DNA damage tolerance (41). Recent work demonstrated that polymerase η is indeed essential for recovery from RS-inducing drugs (42). In addition to the canonical, transcriptional-dependent, function of P53, a growing body of evidence indicates that P53 also executes its function directly at the DNA replication fork (43,44), all together pointing towards a prime function for P53 during S/G2 phase of the cell cycle.

Our live cell imaging data builds upon previous publications which show that cells exit the cell cycle when they encounter severe levels of RS (45,46). It is conceivable that such an exit is important to maintain genomic integrity. Notably, this RS-induced cell cycle exit was delayed in cells with oncogenic RAS (Figure 6C). Several mechanisms can potentially attribute to a prolonged S/G2-phase. Firstly, the speed of S-phase progression correlates with the level of DNA damage. Since HRAS^G12V^-expressing cells experience more RS than wild-type cells during treatment with CHK1i + gemcitabine, this might result in a more profound stalling of DNA replication and subsequently delay the cell cycle exit (34). However, our fiber assays did not show difference in fork speed between cells with and without oncogenic RAS, suggesting that another putative mechanism is in place. This second mechanism involves the P53-P21 pathway, which is at least in part responsible for cell cycle exit by inhibition of CDK2 (45,47,48) and which absence delays cell cycle exit (46). In line with this, it can be hypothesized that the reduced levels of P21 observed in HRAS^G12V^ G2-phase cells contribute to a delayed cell cycle exit from G2.

We have previously shown that cells which exit the cell cycle in G2-phase are not permanently arrested but can re-enter the cell cycle (45). Also after severe drug-induced RS recovery of HRAS-transformed cells occurs, albeit at a lower level (Figure 2E-G). In light of cancer therapy it is of great importance to understand how oncogenic RAS can interfere with this arrest. Reyes and coworkers demonstrated that rare cells which escape a DNA damage-induced cell cycle arrest are characterized by a lower P53 pulse amplitude (49). In such a scenario oncogenic RAS would facilitate cell cycle re-entry by lowering P53 levels. In support of this notion, it was determined that intermediate levels of P21 create a sweet spot for escape from senescence (50). The decision to proliferate or arrest after DNA damage is determined by competing actions of the P53-P21 axis and ERK activity. Besides the inhibitory effect of oncogenic RAS on P53, ERK promotes cell cycle entry by stimulating Cylin D-CDK activity (22). This combined action of oncogenic RAS signaling could tip the balance to cell proliferation at the expense of increased genomic instability. Thus, although RAS-mutant cells accumulate more DNA damage after replication stress, rare cells might be able to escape a G2 arrest. This increases the chance of mutations which can confer drug resistance resulting in tumor relapse.

Besides the pivotal role of P53 in anti-cancer therapy responses, the finding that oncogenic RAS downregulates P53 sheds a new light on tumor evolution. The lack of P53 fulfills a central function in tumorigenesis by evading growth suppressors and resistance to programmed cell death (51). This induces genomic instability which is essential to acquire oncogenic traits. Despite the key role of P53 in tumorigenesis, it is mutated relatively late in the process of malignant transformation (52,53). On this basis, it is tempting to speculate that alternative mechanisms are required to loosen the stringency of the P53-induced checkpoint early during tumor development. In line with this, it is shown that the transcriptional output of wild-type P53 correlates with breast cancer tumorigenesis (54). Moreover, reduced levels of P53 could directly contribute to genomic instability as heterozygous deletion of P53 induces RS (29). In such a scenario, the joined induction of oncogenic RS and P53-checkpoint dampening by activation of oncogenes, such as RAS, would form the basis for malignant transformation.

## Methods

### Key resources

Key resources are listed in Table S1.

### Cell lines and cell line generation

RPE1-hTERT, HEK293T, U2OS and UM-UC-3 cell lines were purchased from ATCC and cultured at 37°C, 5% CO_2_. RPE1-hTERT, U2OS and HEK293T cells were cultured in DMEM, UM-UC-3 cells were cultured in EMEM. All media were supplemented with 10% FBS and 1% pen/strep. All cell lines were regularly tested and confirmed mycoplasma negative.

Gemcitabine, Prexasertib and Ceralasertib were purchased from Selleckchem, Nutlin-3a was purchased from Sigma and used at a final concentration of 4nM, 10nM, 1μM and 1μM respectively, unless stated otherwise.

RPE cell lines harboring the Tet Repressor, FUCCI4 system, fluorescently tagged H2B, 53BP1 and HRAS^G12V^ were created using lentiviral transduction with the third-generation lentiviral packaging system as previously described (55). In brief, HEK293T cells were transfected with 10 μg lentiviral packaging plasmids and 10 μg of the construct of interest using PEI. After 2 hours transfection, medium was washed away, lentivirus containing medium was harvested after 48 hours. RPE cells were transduced with lentivirus containing medium supplemented with 8μg/mL Polybrene for 24 hours.

The lentiviral constructs encoding mKO2-SLBP(18-126) and Clover-Geminin(1-110) were a gift from Michael Lin (Addgene plasmid # 83915; http://n2t.net/addgene:83915; RRID:Addgene_83915, Addgene plasmid # 83914; http://n2t.net/addgene:83914; RRID:Addgene_83914). The plasmid encoding Apple-53BP1trunc was a gift from Ralph Weissleder (Addgene plasmid # 69531; http://n2t.net/addgene:69531; RRID:Addgene_69531). The plasmid encoding pLenti-H2B-iRFP720 was a gift from Carlos Carmona-Fontaine (Addgene plasmid # 128961; http://n2t.net/addgene:128961; RRID:Addgene_128961).

The fluorescent tag of truncated 53BP1 was changed to mTurquoise2 using Gibson Assembly. Cells harboring the Tet repressor and HRAS^G12V^ were selected using blasticidin (10μg/ml, 10 days) and puromycin (1.0μg/ml, 5 days) respectively. Cells harboring fluorescent tagged SLBP, Geminin, 53BP1 and H2B were selected by FACS-sorting.

### Mouse experiments

Animal experiment were approved by the Utrecht University Animal Ethics Committee (approval no. AVD108002016626) and performed according to institutional and national guidelines. For xenograft experiments, 1 million cells in 200 μL basic DMEM were injected in the lower and upper flanks of immunocompromised Rj:NMRI-Foxn1nu/nu mice (Janvier Labs). Doxycycline (2g/kg) was administrated ad libitum in pellets to all injected mice (Bio Services). Tumor size was monitored biweekly, and mice were euthanized when one tumor exceeded the size of one cubic centimeter. Tumors were harvested and stored in paraformaldehyde.

### RNAi transfections

For siRNA experiments, cells were transfected with a final concentration of 10nm siRNA targeting the gene of interesting or a scrambled control using Lipofectamine RNAiMAX according to manufacturers’ instructions (Life Technologies, 13778030). The following siRNAs were used: Dharmacon D-001210-02-05 (Scrambled), LQ-003329-00-0002 (TP53), LQ-019150-00-0002 (RREB1), LQ-006423-00-0005 (CEBPβ), LQ-005089-00-0002 (KLF4), LQ-020067-00-0002 (SMAD3) and LQ-011223-00-0002 (KLF9).

### Microscopy

Live cell imaging experiments and downstream analysis were performed as previously described (45). In brief, 1500 RPE cells were seeded in a CELLview slide (Greiner). Images were acquired at 20-minute time intervals for 48 hours in a humidified chamber on a Nikon A1R-STORM microscope using a 40x LWD objective. Cell tracking and foci analysis was performed in a semi-automated manner using the FIJI plugin Trackmate.

For RPE cells, images for 53BP1 foci analysis of snapshots were acquired on a NIKON Ti-E microscope and foci quantification was performed manually using FIJI.

For immunofluorescence staining, cells were seeded on coverslips, fixed using 4% paraformaldehyde for 20 minutes and permeabilized with 0.1% Triton X-100 for 1 minute at room temperature. Samples were blocked with 5% goat serum and incubated with antibodies prior to mounting on slides. Slides were analyzed on a Leica SP8 confocal microscope using a 20x objective. For analysis, ROIs were determined based on the DAPI signal. For γH2AX staining, foci per cell were determined with a custom-made FIJI script using the Find Maxima function. 53BP1 foci in U2OS cells were quantified manually. For each condition at least 100 cells were quantified. Antibodies used and dilutions are listed in Table S2.

DNA fiber assays were performed as described previously (55). Briefly, cells were pulsed with CldU (25μM) for 20 minutes, washed with PBS and pulsed with IdU (250μM) for 20 minutes. Cells were collected, lysed in spreading buffer (200 mM Tris-HCl pH 7.4, 50 mM EDTA, 0.5% SDS), and spread on a microscope slide under an angle of 15 degrees. Cells were fixed in methanol-acetic acid (3:1) and treated with 2.5 mM HCl for 75 minutes prior to blocking and antibody incubation. Slides were analyzed on a Leica SP8 confocal microscope with 63x objective. Length and type of DNA fibers were analyzed in FIJI using the length measurement tool.

### Flow cytometry

For cell sorting experiments, cells were resuspended in DMEM and filtered with 40 μm cell strainers to remove cell clumps. Cell sorting was performed on a BD FACS Fusion based on fluorescent intensity for mKO2-SLBP and mClover-Geminin. For each condition 100 000 cells were sorted and collected in ice-cold PBS.

### Immunoblotting

For immunoblotting cells were washed with ice-cold PBS and lysed in ice-cold RIPA-buffer (50 nM Tris-HCl pH 7.5, 1 mM EDTA, 150 mM NaCl, 0.25% deoxycholate, 1% NP-40) supplemented with NaF (1 mM), NaV_3_O_4_ (1 mM) and protease inhibitor cocktail (11873580001, Sigma Aldrich) after which samples were subjected to a standard SDS-page immunoblot. Antibodies used and dilutions are listed in Table S2.

### RNA-sequencing

For RNA-sequencing S-phase (Geminin-positive/SLBP-positive) RPE and S/G2-phase (Geminin-positive) UM-UC-3 cells were collected using FACS sorting. For each sample three biological replicates were used, and 2 independent clones of RPE cells harboring the FUCCI4 reporter system were included. RNA was isolated using the QIAGEN RNAeasy kit according to manufacturers’ recommendations. Next, samples were subjected to sequencing and single-end reads were checked for quality, aligned, and counted using the USEQ RNA-seq pipeline (version 2.3.0). Briefly, a quality check with FastQC (version 0.11.4) was performed (56), reads were aligned to the human genome (hg19) with STAR (version 2.4.2a) (57) after which post alignment processing and quality control was done with sambamba (version 0.5.8) (58), Picard-toolkits (version 1.141) and bamMetrics (version 2.1.4) respectively. Final read counts were obtained using HTSeq-count (version 0.6.1) (59). Raw read counts were further analysed with DEseq2 (version 1.28.0) (60) using default analysis parameters and the differential gene expression between groups was assessed as shrunken log2 foldchanges (LFC). Differential expression analysis was performed using R (version 3.6.3) and RStudio Desktop (version 1.3.1093). Pathway analysis was performed on significantly changed genes with a fold change of at least 1.5 with the ToppGene application using KEGG pathways (61). Sequencing data is available on Gene Expression Omnibus under accession number GSE168987.

### Quantitative PCR

RNA isolation (QIAGEN, RNAeasy kit), synthesis of cDNA (Thermo Fisher) and qPCR (Bio-Rad, SYBR green master mix) were performed according to manufacturers’ guidelines. GAPDH and 18S were used as reference genes. Fold changes were calculated using the △△Ct method. Primers used in this manuscript are listed in Table S3.

### Colony formation assay

For colony formation assays cells were plated at low density (250 cells/well) in a 6 well plate. Medium was replenished every 48 hours and cells were fixed after 10 days. Fixation was performed with Acitic acid:Methanol (1:7 vol/vol) for 5 minutes, then cells were washed with PBS and incubated with Crystal Violet staining solution (0.5% Crystal Violet, Sigma-Aldrich) for 2 hours at room temperature. Subsequently, dishes were rinsed with water and air-dried. Number of colonies per well was counted manually, each condition was performed in triplicate.

### Quantification and statistical analysis

Microscopy, immunoblot and quantitative PCR experiments were performed three times unless indicated otherwise. Statistical analysis of qPCR experiments and 53BP1 foci per cell was done by a Kruskal-Wallis test with a post-hoc Dunnett’s test with Benjamin Hochberg correction. * P < 0.05, ** P < 0.01, *** P < 0.001.

## Acknowledgements

We thank Reinier van der Linden and Stefan van der Elst (Hubrecht Institute-KNAW, NL) for assistance with FACS sorting experiments. We thank Esther van ‘t Veld and Richard Wubbolts (Center for Cell Imaging, Utrecht University) for support with live cell imaging experiments. We thank Utrecht Sequencing Facility for providing sequencing service and data. This work is financially supported by the KWF Kankerbestrijding (Dutch Cancer Society, project grant 11941-2018-II) and ZonMW (grant 91116011). Further financial support was provided by research infrastructure grants from Utrecht Life Sciences to the Single Cell Analysis Center and the Center for Cell Imaging. Utrecht Sequencing Facility is subsidized by the University Medical Center Utrecht, Hubrecht Institute, Utrecht University and The Netherlands X-omics Initiative (NWO project 184.034.019).

## Author Contributions

H.A.S. conceived and performed experiments, analyzed data, and wrote the manuscript. E.A.v.L performed experiments and analyzed data. F.M.R. analyzed RNA-sequencing data. A.d.B. provided mentorship and feedback. B.W. conceived and oversaw the study, and wrote the manuscript.

