## Supplemental information and figures for "Oncogenic RAS sensitizes cells to drug-induced replication stress via transcriptional silencing of P53"

### 1. Supplementary Tables

**Table S1. Key resources**

| Reagent or resource | Source | Identifier |
|---|---|---|
| **Antibodies** | | |
| Anti-53BP1 | Novus Biologicals | NB100-304, RRID:AB_10003037 |
| Rabbit monoclonal anti CHK1, Phospho (Ser345) | Cell Signaling Technology | 2348; RRID:AB_331212 |
| Rabbit monoclonal anti CHK1, Phospho (Ser296) | Cell Signaling Technology | 2349; RRID:AB_2080323 |
| Mouse monoclonal anti-CHK1 | Cell Signaling Technology | 2360; RRID:AB_2080320 |
| Rabbit polyclonal anti-P21 | Santa Cruz Biotechnology | sc-471; RRID:AB_632123 |
| Mouse monoclonal anti-P53 | Santa Cruz Biotechnology | sc-126; RRID:AB_628082 |
| Mouse monoclonal anti-gamma-Tubulin | Sigma-Aldrich | T6557; RRID:AB_477584 |
| Rabbit anti-Phospho-Histone H2A.X (Ser139) | Cell Signaling Technology | 2577; RRID:AB_2118010 |
| Rat anti-BrdU | Bio-Rad | OBT0030G RRID:AB_609567 |
| Mouse anti-BrdU | BD Biosciences | 347580 RRID:AB_10015219 |
| **Chemicals** | | |
| Gemcitabine | Selleckchem | S1714 |
| Prexasertib | Selleckchem | S7178 |
| Ceralasertib | Selleckchem | S7693 |
| Nutlin-3a | Sigma-Aldrich | SML0580 |
| DAPI | Sigma-Aldrich | D9542 |
| Protease inhibitor Cocktail | Sigma-Aldrich | 11873580001 |
| Fetal bovine serum | Thermo Fisher | 10500064 |
| DMEM | Thermo Fisher | 41966052 |
| EMEM | LGC | 30-2003 |
| **Critical commercial assays** | | |
| Rneasy Mini Kit for RNA extraction | Qiagen | 74106 |
| Rneasy Micro Kit for RNA extraction | Qiagen | 74004 |
| **Deposited data** | | |

| | | |
|---|---|---|
| RNA sequencing RPE-FUCCI4 $HRAS^{G12V}$ and UM-UC-3 cells | This paper | GEO: GSE168987 |
| **Experimental Models** | | |
| hTert-RPE1 | ATCC | CRL-4000; RRID:CVCL_4388 |
| HEK293T | ATCC | CRL-3216; RRID:CVCL_0063 |
| UM-UC-3 | ATCC | CRL-1749; RRID:CVCL_1783 |
| U2OS | ATCC | HTB-96; RRID:CVCL_0042 |
| mice | Janvier Labs | Rj:NMRI-Foxn1nu/nu mice |
| **Oligonucleotides** | | |
| Primers used for qPCR, see Table S1 | This paper | N/A |
| Scrambled siRNA | Dharmacon | D-001210-02-05 |
| Human P53 siRNA | Dharmacon | LQ-003329-00-0002 |
| Human RREB1 siRNA | Dharmacon | LQ-019150-00-0002 |
| Human CEBPB siRNA | Dharmacon | LQ-006423-00-0005 |
| Human SMAD3 siRNA | Dharmacon | LQ-020067-00-0002 |
| Human KLF4 siRNA | Dharmacon | LQ-005089-00-0002 |
| Human KLF9 siRNA | Dharmacon | LQ-011223-00-0002 |
| **Recombinant DNA** | | |
| pLenti CMV TetR Blast (716-1) | Addgene | Addgene_17492 |
| pLenti CMV $HRAS^{G12V}$ Puro | | N/A |
| Apple-53BP1trunc | Addgene | Addgene_69531 |
| Clover-Geminin(1-110) | Addgene | Addgene_83915 |
| mKO2-SLBP(18-126) | Addgene | Addgene_83914 |
| H2B-iRFP670 | Addgene | Addgene_128961 |
| pMDLg/pRRE lentiviral packaging | Addgene | Addgene_12251 |
| pRSV-Rev lentiviral packaging | Addgene | Addgene_12253 |
| pCMV-VSV-G lentiviral packaging | Addgene | Addgene_8454 |
| **Software and Algorithms** | | |
| FIJI (ImageJ) | https://fiji.sc | RRID:SCR_002285 |
| TrackMate | https://imagej.net/TrackMate | N/A |
| NIS elements | Nikon | RRID:SCR_014329 |
| FlowJo | BD | RRID:SCR_008520 |
| R | https://www.R-project.org/ | N/A |
| Rstudio | https://www.rstudio.com/ | RRID:SCR_000432 |

| | | |
|---|---|---|
| USEQ RNA-seq pipeline | https://github.com/UMCUGenetics/RNASeq | N/A |

**Table S2. Antibodies for immunoblots and immunofluorescence staining**

| *Application* | *Name* | *Company* | *Catalogue number* | *Dilution* |
|---|---|---|---|---|
| *Immunoblots* | CHK1 phospho S296 | Cell Signaling | 2349 | 1:1000 |
| | CHK1 phospho S345 | Cell Signaling | 2348 | 1:1000 |
| | CHK1 total protein | Cell Signaling | 2360 | 1:1000 |
| | ERK1/2 phospho T202/204 | Cell Signaling | 4370 | 1:1000 |
| | ERK1 | Santa Cruz | sc-94 | 1:1000 |
| | γ-H2AX (S139) | Cell Signaling | 2577 | 1:1000 |
| | P21 | Santa Cruz | sc-471 (M-19) | 1:1000 |
| | P53 | Santa Cruz | sc-126 | 1:1000 |
| | γ-Tubulin | Sigma Aldrich | T6557 (GTU-88) | 1:1000 |
| *Immunofluorescence* | γ-H2AX (S139) | Cell Signaling | 2577 | 1:200 |
| | 53BP1 | Novus Biologicals | NB100-304 | 1:2000 |
| | Rat anti-BrdU | Bio-Rad | OBT0030G | 1:100 |
| | Mouse anti-BrdU | BD Biosciences | 347580 | 1:100 |

**Table S3. qPCR primers**

| *Gene* | *Forward primer (5'-3')* | *Reverse primer (3'-5')* |
|---|---|---|
| *BTG2* | GGCTTAAGGTCTTCAGCGGG | TGTGGTTGATGCGAATGCAG |
| *BBC3* | GAGCAGGGCAGGAAGTAACA | CACAAATCTGGCAGGGGACC |
| *CDKN1A* | CTCTAAGGTTGGGCAGGGTGACC | CAGAGGGGGGTATCAAGAGCCAG |
| *CEBPβ* | TTTGTCCAAACCAACCGCAC | GCATCAACTTCGAAACCGGC |
| *GAPDH* | CTCTGCTCCTCCTGTTCG | GCCCAATACGACCAAATCC |
| *HRAS* | GGAAGCAGGTGGTCATTGAT | ATGGCAAACACACACAGGAA |
| *KLF4* | CATCTCAAGGCACACCTGCGAA | TCGGTCGCATTTTTGGCACTGG |
| *KLF9* | GCCGCCTACATGGACTTCG | GGATGGGTCGGTACTTGTTCA |
| *MDM2* | CGAGCTTGGCTGCTTCTGG | GTACGCACTAATCCGGGGAG |
| *PCNA* | GGTTACTGAGGGCGAGAAGC | GCTGAGACTTGCGTAAGGGA |
| *POLH* | ACTGGCACAAGTTCGTGAGT | CATCAATGCTGGCACGTTCA |
| *RREB1* | CATGCTCACACACACTGGTC | CGTGAGGTGAGGTCTAGCAC |
| *RSP18* | AGTTCCAGCATATTTTGCGAG | CTCTTGGTGAGGTCAATGTC |
| *SMAD3* | CATGGACGCAGGTTCTCCAA | GGCTCGCAGTAGGTAACTGG |
| *TP53* | GTTCCGAGAGCTGAATGAGG | TCTGAGTCAGGCCCTTCTGT |
| *XPC* | TTGTCGTGGAGAAGCGGTCTAC | CTTCTCCAAGCCTCACCACTCT |

### 2. Supplementary Figure Legends

**Figure S1. Related to Figure 1**

**A** Quantitative PCR of HRAS 24 hours after doxycycline administration in RPE cells with inducible $HRAS^{G12V}$.

**B** Immunoblot showing elevated levels of phosphorylated ERK1/2 in cells with oncogenic HRAS.

**C** Dot plot showing the length of cell cycle phases in cells with or without oncogenic RAS measured using live cell imaging. 100 cells per condition were analyzed. Statistical differences were evaluated using a Kruskal-Wallis test and post-hoc Dunnett's test.

**Figure S2. Related to Figure 1**

**A** Quantification of different types of DNA replication forks observed in a DNA fiber assay in the absence of drugs. Bars represent mean ± s.e.m. of 3 independent experiments, per experiment at least 250 fibers were analyzed. Statistical differences were evaluated using a Kruskal-Wallis test and post-hoc Dunnett's test.

**B** Dot plot showing the replication fork speed of individual DNA tracks measured using a DNA fiber assay. Per condition at least 100 fibers were measured.

**C** Immunoblot showing levels of phosphorylated CHK1 in the absence and presence of oncogenic RAS.

**D** Bar chart showing the number of $\gamma$H2AX foci, as read out for DNA damage, per cell in the absence or presence of oncogenic RAS. At least 100 cells per condition were analyzed. Statistical differences were evaluated with a Chi-Square test.

**Figure S3. Related to Figure 2**

**A** Dot plot showing the replication fork speed of individual DNA tracks, before and 16 hours after treatment with CHK1i + gemcitabine, measured using a DNA fiber assay. Per condition at least 250 fibers were measured. Statistical differences were evaluated using a Student's t-test.

**B** Cell proliferation, as measured by cell counting, of RPE WT or $HRAS^{G12V}$ cells with or without doxycycline and CHK1i + gemcitabine.

**C** Quantitative PCR of HRAS 24 hours after doxycycline administration in U2OS cells with inducible $HRAS^{G12V}$ in the absence and presence of CHK1i + gemcitabine.

**D** Quantification of immunofluorescence staining of 53BP1 in U2OS cells. Bar chart shows the number of 53BP1 foci, as read out for RS, in individual cells in the absence or presence of oncogenic RAS. At least 100 cells per condition were analyzed. Statistical differences were evaluated with a Chi-Square test corrected for multiple comparisons.

**E** Same as D, but now in the presence of CHK1i + gemcitabine.

**F** Quantification of colony formation assay in RPE WT and RPE $HRAS^{G12V}$ cells with or without doxycycline, 10 days after cell plating. Bars represent mean ± s.e.m.

**Figure S4. Related to Figure 3**

**A** Sorting strategy to collect G1, S and G2-phase cells based on the FUCCI4 system.

**B** Pathway analysis of differentially expressed, up or downregulated, genes in S-phase cells with (dox) and without (veh) $HRAS^{G12V}$. Significantly changed genes with a fold change of at least 1.5 were selected for analysis. Red dotted line indicates *P* value of 0.05.
**C** Venn diagrams of significantly changed genes, with a fold change of at least 1.5 before and after treatment with CHK1i + gemcitabine. RPE $HRAS^{G12V}$ S-phase cells with or without doxycycline and UM-UC-3 S/G2-phase cells were analyzed.
**D** Cell proliferation, as measured by cell counting, of RPE WT and UM-UC-3 cells with or without CHK1i + gemcitabine.
**E** Quantitative PCR of *TP53* 24 hours after doxycycline administration in U2OS cells with inducible $HRAS^{G12V}$ in the absence and presence of CHK1i + gemcitabine. Bars represent mean ± s.e.m. of 2 independent experiments. Statistical differences were evaluated using a Kruskal-Wallis test and post-hoc Dunnett's test.

**Figure S5. Related to Figure 4**
**A** Quantitative PCR of siRNA target (*RREB1*), *TP53* and P53 targets 24 hours after transfection of RPE $HRAS^{G12V}$ cells in the presence or absence of doxycycline. Bars represent mean ± s.e.m. of 2 independent experiments. Statistical differences were evaluated using a Kruskal-Wallis test and post-hoc Dunnett's test.
**B** Table with putative P53 transcriptional regulators selected based on RNA-sequencing data. Fold changes, on a linear scale, between RPE cells with and without $HRAS^{G12V}$ in the absence (cntr) and presence of RS-inducing drugs (CHK1 + gemc) as detected using RNA-sequencing are shown.
**C** Same as A, but know with siRNA targeting *CEBP–beta*.
**D** Same as A, but know with siRNA targeting *KLF4*.
**E** Immunoblot showing efficient knock down of SMAD3 in RPE cells 24 hours after transfection.
**F** Quantitative PCR of siRNA target (*SMAD3*), *TP53* and P53 targets 24 hours after transfection of RPE WT cells in the presence or absence of CHK1i + gemcitabine. Bars represent mean ± s.e.m. of 2 independent experiments. Statistical differences were evaluated using a Kruskal-Wallis test and post-hoc Dunnett's test.
**G** Same as F, but know with siRNA targeting *KLF9*.

**Figure S6. Related to Figure 5 and 6**
**A** Quantification of live cell imaging data showing the average number of 53BP1 foci per image per cell in S/G2-phase. Cells were separated based on their fate as shown in Figure 6B.
**B** Quantitative PCR showing increase in P53 target gene expression when RPE cells are treated with Nutlin-3a in addition to CHK1i + gemcitabine. Cells were harvested 24 hours after treatment. Bars represent mean ± s.e.m. of 2 independent experiments. Statistical differences were evaluated using a Kruskal-Wallis test and post-hoc Dunnett's test.
**C** Heatmap showing cell cycle progression and 53B1 foci in RPE WT cells which were in S-phase at the start of imaging. Cells were traced until they completed mitosis or exited the cell cycle.
**D** Same as C, but now with RPE $HRAS^{G12V}$ dox cells.

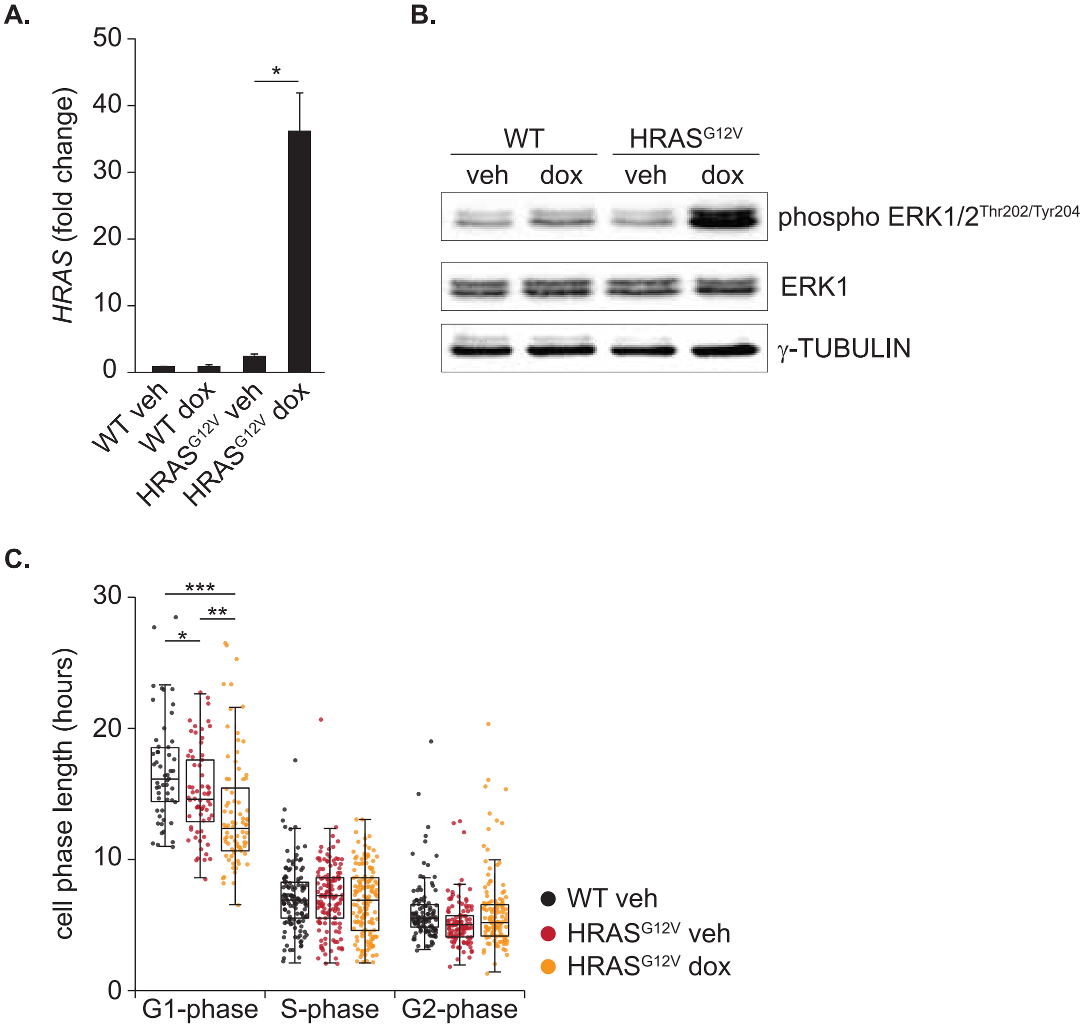


**Supplemental Figure 1, related to Figure 1**

A.

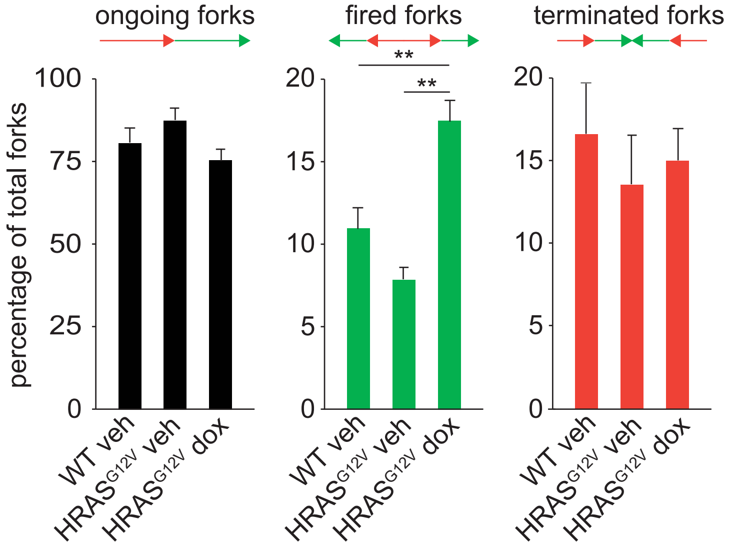


B.

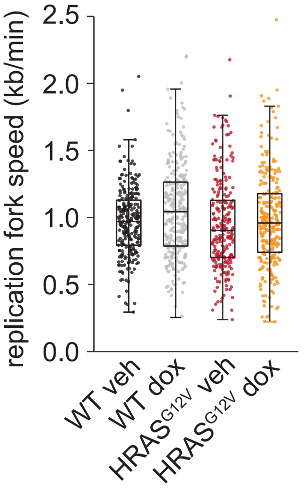


C.

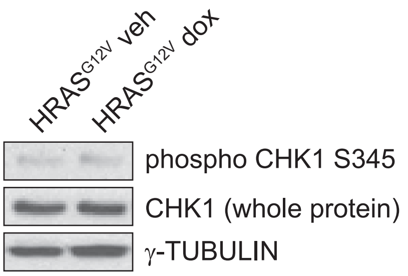


D.

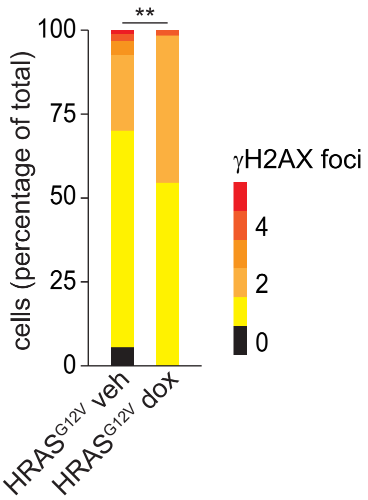


**Supplemental Figure 2, related to Figure 1**

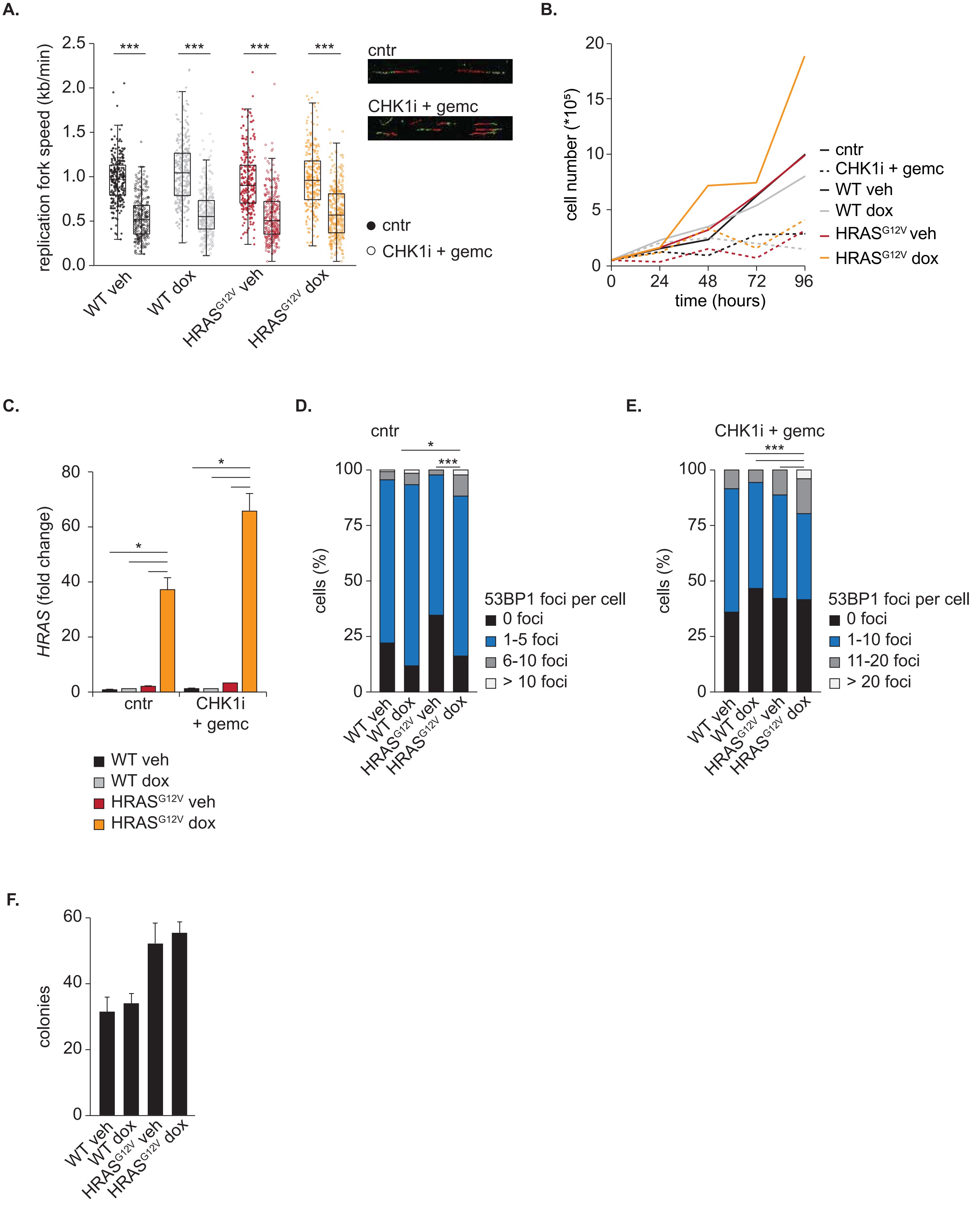


**Supplemental Figure 3, related to Figure 2**

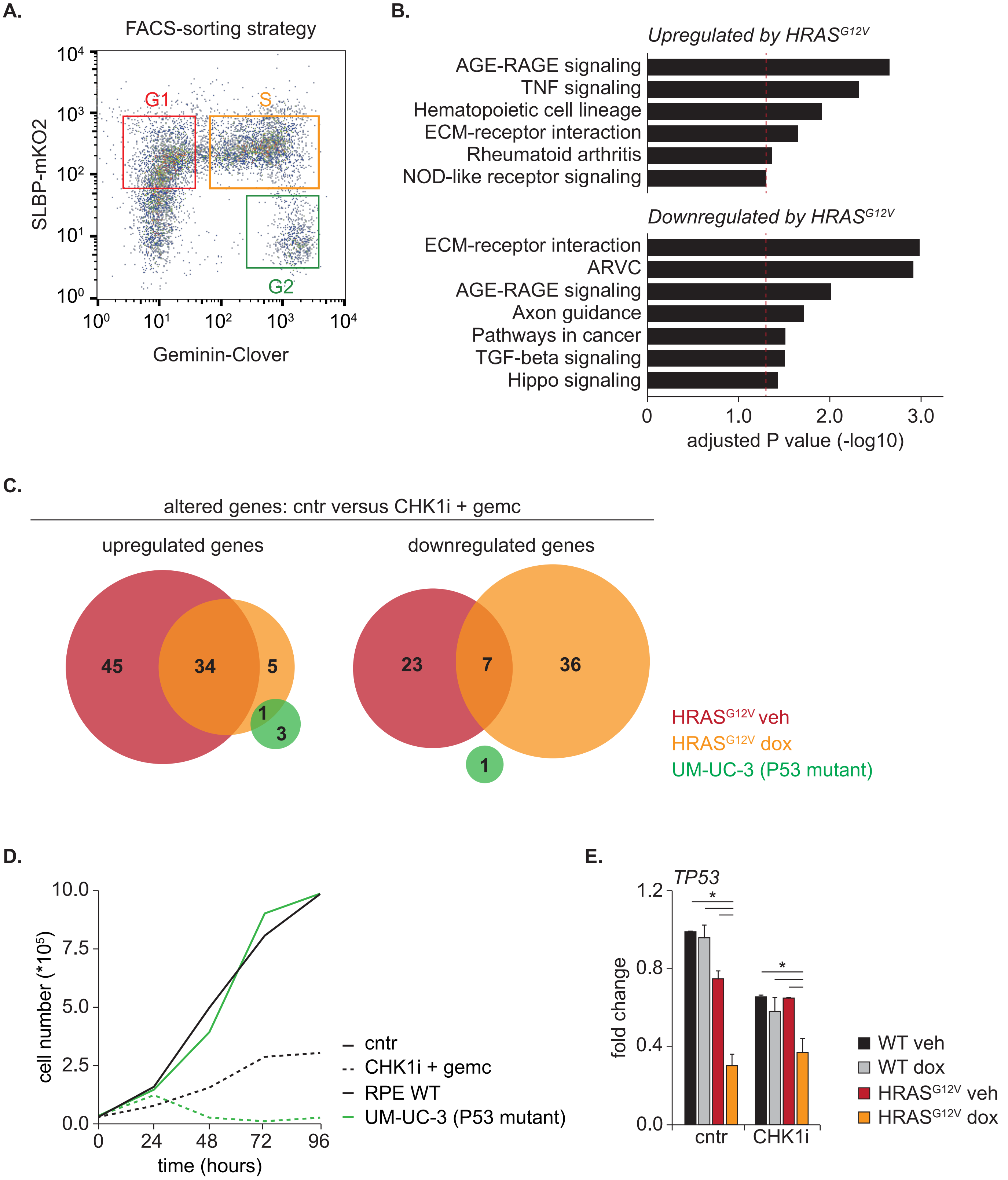


**Supplemental Figure 4, related to Figure 3**

A.

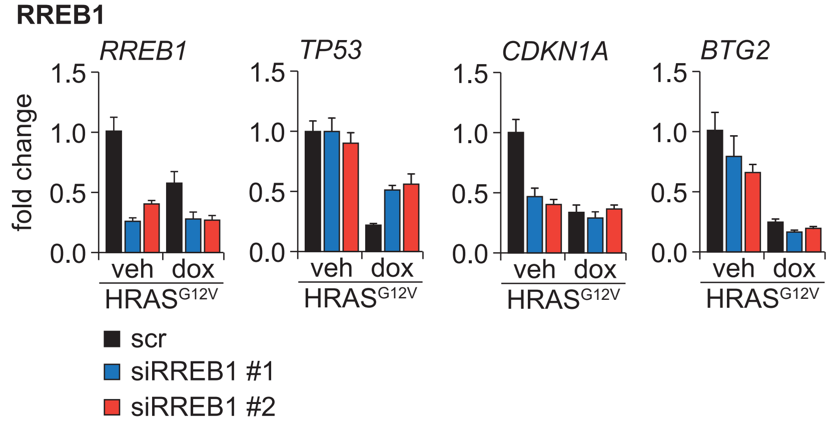


B.

| putative transcriptional regulators of P53 | | | |
|---|---|---|---|
| gene | function | fold change cntr | fold change CHK1i + gemc |
| *CEBPβ* | repressor | 2.55 | 3.58 |
| *KLF4* | repressor | 1.68 | 1.54 |
| *SMAD3* | activator | - 1.61 | - 1.60 |
| *KLF9* | activator | - 2.78 | - 2.97 |

C.

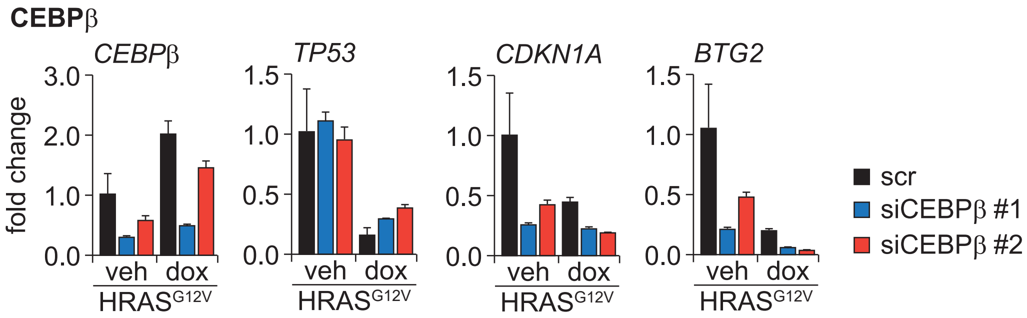


D.

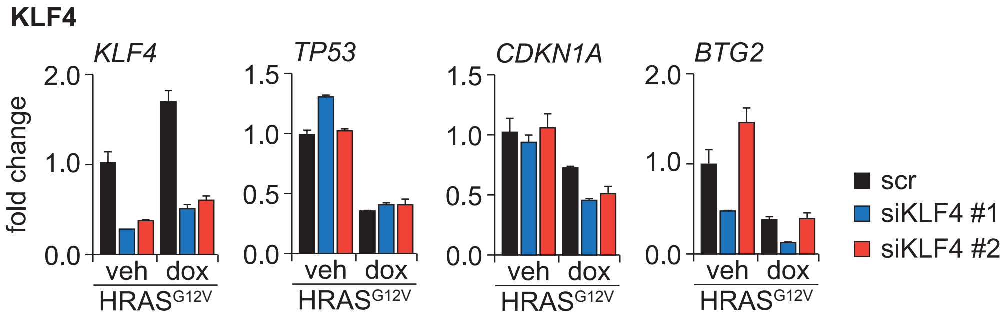


E.

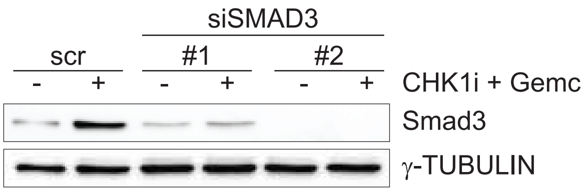


F.

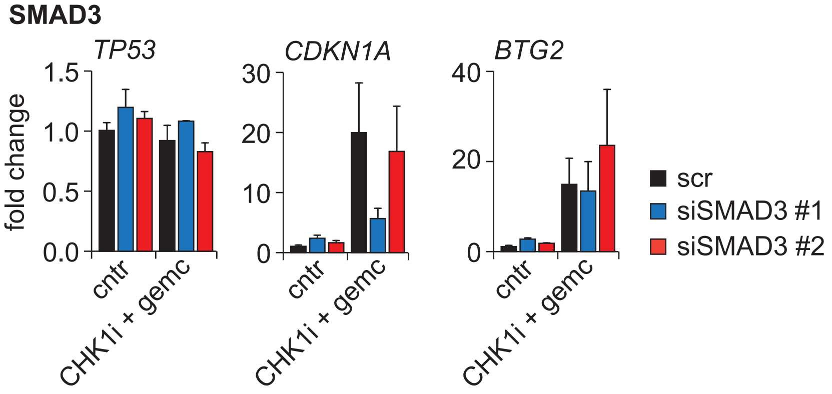


G.

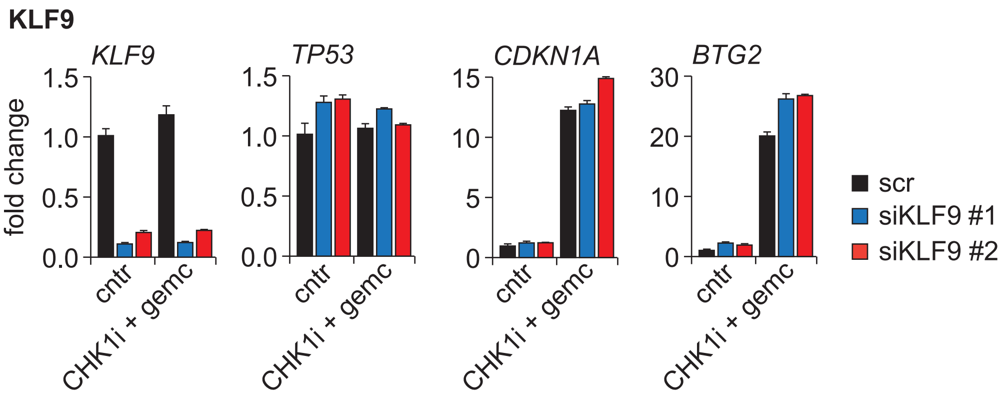


**Supplemental Figure 5, related to Figure 4**

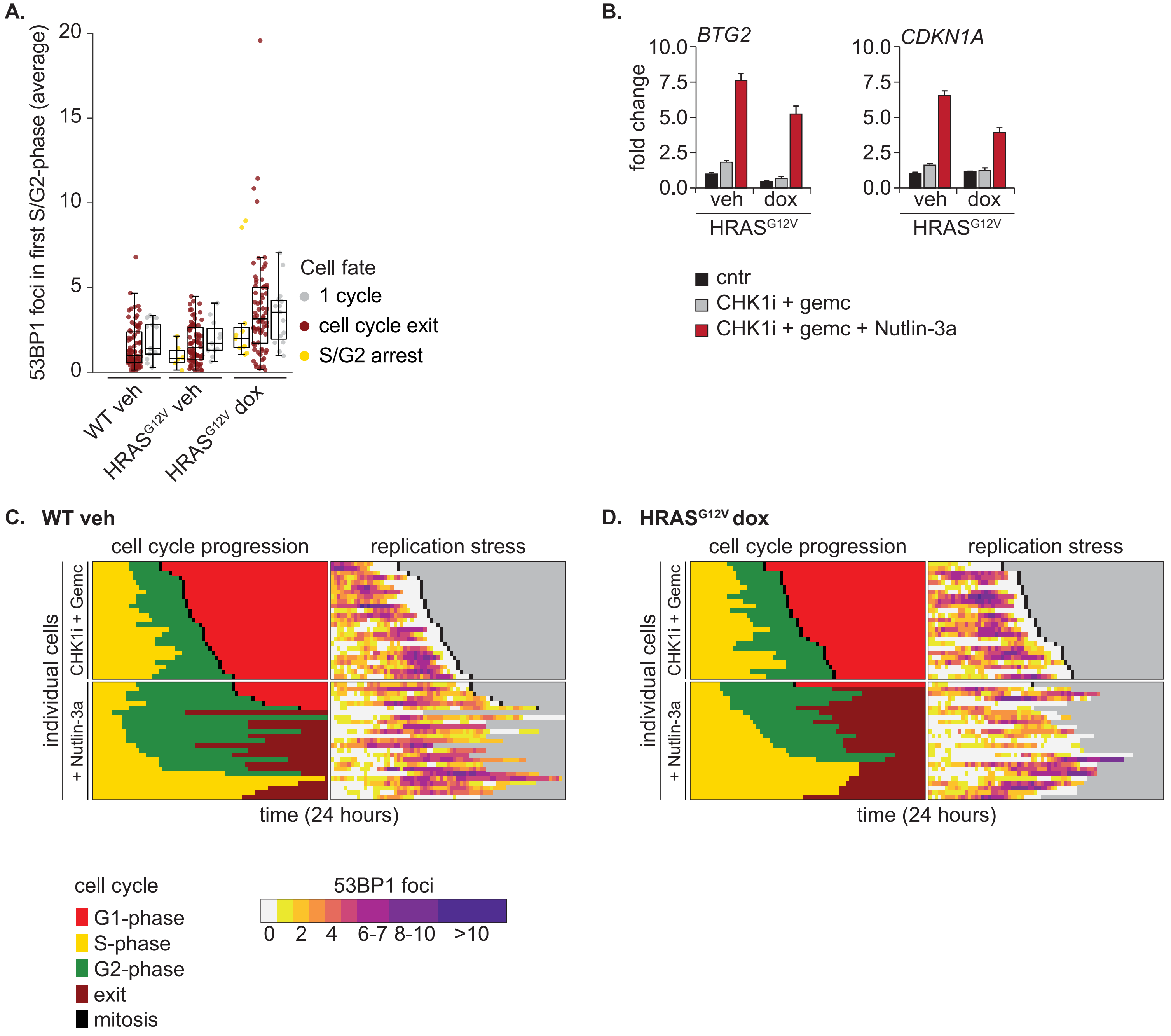


**Supplemental Figure 6, related to Figure 5 and 6**
